# Prestimulus alpha phase modulates visual temporal integration

**DOI:** 10.1101/2024.01.29.577737

**Authors:** Michelle Johannknecht, Alfons Schnitzler, Joachim Lange

## Abstract

When presented shortly after another, discrete pictures are naturally perceived as continuous. The neuronal mechanism underlying such continuous or discrete perception are not well understood. While continuous alpha oscillations are a candidate for orchestrating such neuronal mechanisms, recent evidence is mixed. In this study, we investigated the influence of prestimulus alpha oscillation on visual temporal perception. Specifically, we were interested whether prestimulus alpha phase modulates neuronal and perceptual processes underlying discrete or continuous perception. Participant had to report the location of a missing object in a visual temporal integration task, while simultaneously MEG data was recorded. Using source reconstruction, we evaluated local phase effects by contrasting phase angle values between correctly and incorrectly integrated trials. Our results show a phase opposition cluster between - 0.8 to - 0.5 s (relative to stimulus presentation) and between 6 - 20 Hz. These momentary phase angle values were correlated with behavioural performance and event related potential amplitude. There was no evidence that frequency defined a window of temporal integration.

**Significance Statement:** In light with the current debate if our visual perception is a rhythmic or discrete process, we give new insight to this debate. We investigated potential underling mechanism defining potential rhythmic perception and highlight the complexity of this process. This will help us further understand how our brain operates and processes incoming unimodal visual stimuli. In a visual temporal integration task, we were able to show that the incoming information were processed in rhythmic fashion. Our data supports the idea that the phase of prestimulus alpha oscillation modulates poststimulus visual processing by defining good and less good phases for early visual processes. We were not able to show that prestimulus alpha oscillation defines windows were two visual stimuli are integrated into one single event.

## Introduction

Although our everyday experience implies that perception is a seamless process, the human perceptual system is limited to accurately detect and process incoming information. These limitations affect e.g., the spatial but also the temporal resolution of perception. For example, a series of discrete stimuli will be perceived as a continuous, seamless flow of information, if the presentation duration is faster than the temporal resolution of the visual system – like in movies.

Research in the last decades has highlighted a functional role of neuronal oscillations for perception. Especially prestimulus alpha oscillations have been shown to be a relevant factor for perception near threshold. For example, recent studies demonstrated, the power of alpha oscillations influences perception in near-threshold detection tasks (van Dijk et al., 2008; Lange et al., 2013; Baumgarten et al., 2016; Limbach and Corballis, 2016; Iemi et al., 2017; Benwell et al., 2022).

In addition, evidence showed that the momentary phase of prestimulus oscillations influences the detectability of stimuli in near-threshold detection task (Busch et al., 2009; Mathewson et al., 2009; Busch and VanRullen, 2010; Dugue et al., 2011; Landau and Fries, 2012). While most of the evidence for a potential role of phase for perception stems from detection tasks, few studies also reported an influence of phase on temporal perception (Baumgarten et al., 2015; Milton and Pleydell-Pearce, 2016; Ikumi et al., 2019).

Despite the accumulating evidence for a functional role of alpha phase for visual perception, there are still unresolved questions and remaining concerns. For instant, while some studies reported an influence of alpha phase on perception, other studies reported null results (summarized by Keitel et al., 2022). Moreover, inconsistencies in the reported frequency ranges in similar tasks raise the question of validity of these findings (Merholz et al., 2022). Finally, the functional role of prestimulus phase remains unclear. Prestimulus phase might modulate sensory processing by modulating poststimulus evoked potentials (Busch and VanRullen, 2010; Milton and Pleydell-Pearce, 2016). Other studies propose that the phase effects may indicate that specific frequency bands are relevant for perceptual integration or segregation of uni- or multimodal stimuli (Baumgarten et al., 2015; Wutz et al., 2018; Ikumi et al., 2019).

While most studies focused on the role of phase for detection, we investigated the role of phase for temporal integration of visual stimuli. Previous studies investigated the functional frequency range for visual integration (Wutz et al., 2018). We expand these finding by focusing on prestimulus phase. In addition, we aimed to determine whether the prestimulus phase influences temporal integration mechanisms by modulating early poststimulus sensory processing or whether the frequency of prestimulus phase effects determines temporal integration windows. To investigate these hypotheses, participants performed a visual temporal integration study (Wutz et al., 2016), while we simultaneously recorded their neural activity using MEG. We analysed whether the phase of neuronal oscillations correlated with successful integration of the visual stimuli and poststimulus sensory processing.

## Material and Methods

### Participant and ethical information

We recruited 25 participants (17 female, mean age 25.4 ± 5.1 years [SD]) for this study. We chose the number of participants based on previous paper using the same task (Wutz et al., 2016, 2018). The participants provided written informed consent in line with the Declaration of Helsinki and the study was approved by the Ethical Committee of the Medical Faculty, Heinrich Heine University. No participant reported to have any neurological or psychological disorder. All participants had normal or corrected-to-normal vision. Participants received 10€ per hour for participation.

### Stimuli and Task

We adapted the stimuli and task design from Wutz et al. (2016). Each trial started with a randomized prestimulus period between 1200 – 1600 ms with a central fixation cross, followed by the presentation of the stimulus (Fig. 1). A stimulus consisted of two images presented for 16 ms, each. Both images were separated by individual stimulus onset asynchronies (SOAs, see below for details on the SOA). Each image showed seven full annuli and one-half of an annulus presented on pseudo-random positions on a 4 x 4 grid. When both images are integrated, annuli fill out each position, except one. A post-stimulus period followed the stimuli, with a random duration between 600 – 1200 ms with only a fixation cross. Finally, an instruction text appeared which prompted participants’ responses. Participants’ task was to report the empty position. Importantly, participants could report the position only if they temporally integrated both images. Participants reported the position of the missing element by responding twice: The first response was to report the number of the row of the missing element and the second response to report the column number within the 4x4 grid. Participants responded with their right hand via button press. When they responded too early (before presentation of the instructions) or too late (>2000 ms after presentation of instructions), they received feedback accordingly and the trials was appended to the end of the block.

**Figure 1:**
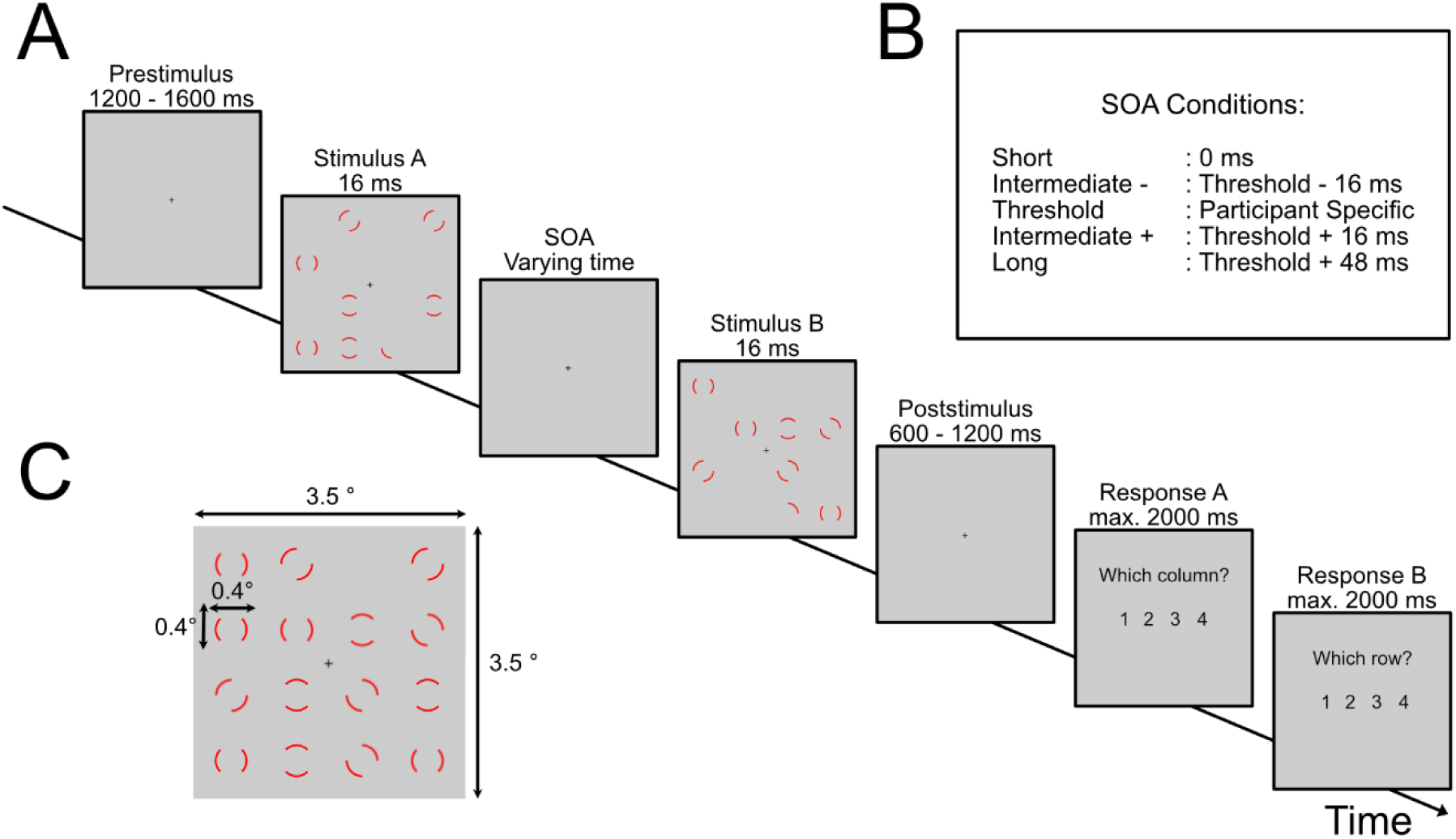
Paradigm. (A) The experiment started with a prestimulus phase (randomly between 1200-1600 ms) where only a fixation cross was presented. Afterwards the first stimulus was presented for 16 ms, followed by a stimulus offset asynchrony (SOA) and the second stimulus. After a poststimulus period (randomly between 600-1200 ms), participants reported the position of the empty location. (B) The different SOA conditions used in the experiment. The threshold SOA was individually determined to achieve 50 % accuracy. (C) An example of a full integration of the two stimuli.

We projected the stimuli on a translucence screen using a projector located outside the magnetically shielded room (Panasonic, PT-DW700E; 60 Hz refresh rate) and a mirror system. The screen was 140 cm away from the participant. Grid dimensions were 8 x 8 cm (i.e., visual angle of 3.5° x 3.5°, each annuli was 1 x 1 cm in size; annuli were evenly spaced on the grid). A training session of ∼2-5 min preceded the experiment.

Before the main experiment, we used a staircase method to measure the performance threshold of 50 % performance accuracy. The staircase started with a fixed SOA of 26 ms. The SOA increased when participants answered twice in a row correct or decreased when they answered twice incorrect. The step size of the increase/decrease was 16 ms (e.g. one frame). When performance was stable at 50% accuracy over the last 20 trials (i.e., the variance of SOAs was < .5), the staircase terminated and the current SOA was taken as the threshold SOA. The staircase was interleaved with SOAs randomly picked from the predefined SOA distribution (starting at 0 ms and increased in steps of 16 ms up to 144 ms). These random SOAs were not included in the computation of the threshold SOA. The threshold SOA is the temporal offset between the two images the participant needed to integrate both images in 50 % of trials. We used additional SOA conditions (0 ms [i.e., both images presented directly after each other], threshold SOA ± 16 ms, threshold SOA + 48 ms) to monitor participants’ behaviour (Baumgarten et al., 2015, 2017). One block consisted of two repetitions of the long (threshold SOA ± 48 ms) and short SOA (0 ms) each, four repetitions for each of the intermediate conditions, and 15 repetitions of the threshold SOA, all stimuli presented in pseudo-randomized order. The entire experiment consisted of 15 blocks and lasted between 30 min and 45 min. After every 100 trials, the participant could take a self-paced break. We used the software Presentation (Neurobehavioral System, Albany NY, USA) to control the experiment.

### MEG recording

During the experiment, we recorded the electromagnetic signal of the brain using a 306 channel Magnetoencephalography (MEG) system (MEGIN, Finland), with a sampling rate of 1000 Hz. The electrooculogram (EOG) was simultaneously recorded by placing electrodes above and below the left eye and on the right and left temple. The head position inside the MEG was co-registered using four head position indicator (HPI) coils. We placed the coils behind the left and right ear and on the right and left forehead. We digitized the coil positions, anatomical landmarks (nasion, left and right periauricular points) and ∼50-100 additional points from the head using the Polhemus digitizer (Fastrak, Polhemus, Vermont, USA).

### MRI recording

For each subject, we recorded a structural magnetic resonance image, with a 3-T MRI scanner (Siemens, Erlangen, Germany). Afterwards, MEG data was aligned offline with the MRI using anatomical landmarks (nasion, left and right pre-auricular points).

### Behavioural analysis

We analysed the behavioural data on group level by calculating the fractional accuracy for each SOA condition and subject. We averaged the individual fractional accuracies across participants. To test for statistical difference in accuracy per SOA, we performed a one-way ANOVA and post hoc Tukey-Kramer tests for pairwise comparisons between SOAs. See also table S1 for an overview of the statistical tests performed.

### Preprocessing of MEG data

For the analysis of MEG data, we used MATLAB (R2019b) and the analysis toolbox FieldTrip (Version 20210825; Oostenveld et al., 2011). First, we divided the continuous MEG data into trials starting with the presentation of the fixation cross and ending with the presentation of the response instructions. We removed jump artefacts, eye movement and muscle movement, using a semi-automatic approach implemented in FieldTrip. We applied band-stop filters between 49 Hz to 51 Hz, 99 Hz to 101 Hz and 149 Hz to 151 Hz, to remove power line and a band-pass filter between 2 and 200 Hz to the data. Next, we visually inspected the data to remove noisy channels and trials. Finally, we used independent component analysis (ICA) to remove undetected noise or artefacts. To speed up ICA, we resampled the data to 150 Hz.

### Source projection of MEG data

Source grid models were computed by applying a regular spaced 3D grid with 5 mm resolution to the Montreal Neurological Institute (MNI) template brain provided by the FieldTrip toolbox. We computed individual grids by nonlinearly warping the individual structural MRI on the MNI MRI, after which we applied the inverse of this warp to the MNI template grid, which resulted in the individual warped virtual sensor grid.

Next, we calculated spatial filters in the time domain by computing a lead field matrix for the reconstructed warped visual sensor grid (Nolte 2003), and using data between -1 s and -0.5 s. We calculated spatial filters using the linear constrained minimum variance beamforming approach (Van Veen 1997) and projected the sensor time-series MEG data through these filters. This projection resulted in time-series data for each grid point. We restricted from here further analyses to regions of interest (ROI), defined by the AAL atlas (Tzourio-Mazoyer et al., 2002) implemented in FieldTrip. We expected to see effects in early visual areas and thus restricted the ROI to these areas. We included the following regions (left and right hemisphere): *Calcarine fissure, Precuneus, Lingual gyrus, Cuneus, Superior occipital lobe, Middle occipital lobe, Inferior occipital lobe, Superior parietal gyrus, Inferior parietal gyrus*.

### Calculating phase opposition sum and surrogate data

To analyse whether the phase of neuronal oscillations influences temporal integration, we included only threshold SOA trials. We sorted the trials into three different groups: “correct” (including only trials where the coordinates of the missing element were correctly reported), “incorrect” (trials where the coordinates were not correctly reported), and the combination of correct and incorrect, which will be labelled “all”.

We analysed phase differences between correct and incorrect trials using the phase opposite sum (POS, VanRullen, 2016). To this end, we calculated for each grid point and trial a Fast Fourier Transformation (FFT) for frequencies between 2 Hz to 30 Hz in steps of 2 Hz. The time window of interest was the prestimulus period between -1 s to 0 s (onset of stimulus). We used a sliding window of 0.3 s length, moved in steps of 0.05 s. Prior to FFT, we multiplied the data with a single Hanning taper. Next, we calculated for each condition the inter trial coherence (ITC), which is a measure for phase consistency across trials (VanRullen, 2016).

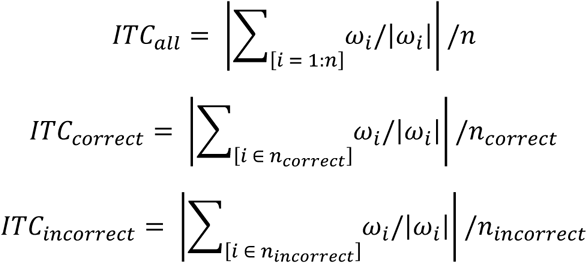

N refers to the number of trials in each group and ω to the angle at trial i. Using the ITC values, we calculated the phase opposition sum (POS).

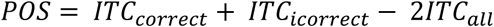

The POS value was calculated for each time-frequency-grid point sample independently. POS values are bound between zero and two, zero indicating that the phase values between correct and incorrect are identical and positive values indicate that phase values are different between correct and incorrect. Since trial number biases ITC, we picked randomly a subset of trials from the group with the higher trial count to equal the number of trials with the group containing fewer trials and then computed ITC and POS values. The average trial number for correct trials was 87.4 (± 34.4 STD) and for the incorrect trials 81.2 (± 25.1 STD). We repeated this step 100 times, resulting in 100 POS values per time-frequency-grid point sample. We calculated the median of POS values per subject and time-frequency-grid point sample.

These observed POS values were statistically tested against a null distribution of surrogate data. We created surrogate data by shuffling the trials from group correct and incorrect, while accounting for equal trial count. We selected 100 times randomly trials and calculated the mean POS values. POS values were calculated as described above. We calculated for each participant 100 mean POS values for each time-frequency-grid point.

To test for group level effects of the POS effect, we used a cluster-based permutation approach (Oostenveld et al., 2011). We first normalized the individual POS values by subtracting the mean and dividing by the standard deviation of the surrogate data. Then, we statistically compared observed and surrogate data across subjects by means of a dependent-samples t-test. We used the Monte Carlo method with 1000 repetition. This step was done independently for each time-frequency-grid point sample. Next, we combined data points with a p-values < 0.05 which were adjacent in time, frequency, and space to a cluster. We pre-defined a neighbour structure for the data, which contains the positions of the channel and its relative neighbours and restricted the neighbours to be maximum 0.6 cm away. Minimum cluster size was set to two.

To obtain the surrogate cluster distribution, we calculated p-values for each time-frequency-grid point and repetition, by testing every surrogate value against the other surrogate values of that time-frequency-grid point. We defined clusters with the same parameters as described for the observed data. This process was repeated 1000 times, always selecting the largest surrogate cluster found. This resulted surrogate cluster distribution was used to test the observed data against it (see table S1).

In an additional post hoc analysis we investigated if power had a systematic influence on our reported phase effects (see Results). We median split the threshold SOA trials into low and high power trials and calculated the POS values as described above. We averaged the POS values for each participant over the channel-time-frequency points where we found phase effects. We applied a t-test to test for systematic differences between phase values between high and low power trials. Phase dependent behavioural performance

### Phase dependent Behaviour

The POS analysis provides information on whether the phase of neuronal oscillations differs between correct and incorrect trials. For a more detailed understanding of the relationship between prestimulus phase and performance accuracy, we calculated the participants’ fractal accuracy for different phase bins. To this end, we first determined in each participant the time-frequency sample with the highest POS value within the grid points belonging to the significant cluster in the POS analysis (Baumgarten et al., 2015). For this time-frequency sample, we computed the momentary phase for each threshold SOA trial (see above for the parameters). Next, we binned the phase values from -π to +π in steps of 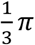. For each of these six phase bins, we calculated the individual performance averaged across all trials. We normalized individual performance by subtracting the mean of all bins from each bin. Next, we aligned the bins so that the bin with the highest performance was aligned to bin 0. Finally, we averaged the data per bin across participants. To test for statistical significance, we performed a one-way repeated ANOVA (see table S1), while bin 0 was excluded from the statistics, and post hoc Tukey-Kramer tests for pairwise comparisons between phase bins. The average trial count per bin was for bin one 27.7 (± 7.7 SD), bin two 28.2 (± 8.2 SD), bin three 26.0 (± 7.4 SD), bin four 29.6 (± 8.6 SD), bin five 27.2 (± 7.4 SD) and bin six 28.5 (± 6.7 SD).

### Phase dependent ERF

We investigated whether prestimulus phase affects visual stimulus processing. To this end, we used an early visual components of the event related field (ERF) – the N170/N1 component - as a measure of stimulus processing (e.g., Mazaheri and Picton, 2005; Han et al., 2015). The individual latency of the component was based on a data driven approach. We took the first component in our data, which had a negative deflection at around 170 ms., hence we called it N170 component. First, we determined 100 grid points showing the highest ERF values. To this end, we calculated for each grid point t-values for the ERF by applying a dependent sample t-test between post-stimulus ERF and prestimulus ERF (averaged across -0.8 s to -0.5 s) and selected 100 grid points showing the highest absolute t value averaged across 170 ms ± 25 ms.. The N170 component was therefore based on an average of times points, this was done to account for varying time point between subjects. Next, from these 100 selected channels, we computed for each participant the momentary phase for the threshold SOA trials. We binned phase as described above and re-calculated the ERF t values, this time only for the trials within each bin. We averaged the ERF t value across all 100 grid points for each participant and aligned the highest value to bin zero. Finally, we averaged the t-values per bin over participants and performed a one-way repeated measures ANOVA, excluding bin with phase 0, and post hoc Tukey-Kramer tests for pairwise comparisons between phase bins (see table S1). The average trial count for the six bin was for bin one 26.5 (± 8.0 SD), bin two 27.5 (± 6.3 SD), bin three 29.1 (± 8.6 SD), bin four 30.2 (± 8.3 SD), bin five 26.3 (± 7.5 SD) and bin six 27.5 (± 7.0 SD).

### Correlation of peak frequency with threshold SOA

Previous studies reported a correlation between participants’ perceptual temporal resolution and the peak frequency of neuronal oscillations (e.g., Samaha et al., 2015). Based on these findings, we tested if the individual peak frequency correlates with the SOA derived from the staircase and the theoretical fitted SOA. For the theoretical fitted SOA, we fitted a sigmoid function onto the individual behavioural data using the Palamedes toolbox for MATLAB (Prins and Kingdom, 2018). From the fitted function, we computed the SOA for which the participant reached 50% correct responses. This SOA, we call the fitted threshold SOA. We used this second approach to account for potential noise in the initial threshold SOA estimation. We estimated the peak frequency for each participant not based on power values but on the highest POS value. To this end, we selected within the significant cluster in the POS analysis the frequency of the highest POS value per participant (using the Matlab function findpeaks.m). Finally, we correlated (Pearson’s correlation) the estimated individual peak frequencies with the individual threshold SOA estimated during the staircase and with the fitted SOA (fitted SOA) (see table S1). Additionally, we fitted a general linear model onto the data for visualization. We are aware that this is not a conventional method to investigate the influence of peak frequency on the temporal integration window (Cecere et al., 2015; Samaha and Postle, 2015; Baumgarten et al., 2017; Tarasi and Romei, 2023). Therefore, we ran a complementary analysis to detect power peaks and correlated them with the staircase and fitted threshold SOA. The prestimulus power spectrum was calculated using a single Hanning window with a frequency resolution of 2 Hz, between 2 and 30 Hz. With the FOOOF toolbox (Version 1.1.0) (Donoghue et al., 2020), we corrected for the 1/f component. We determined for each virtual channel the frequency with the highest power using findpeak.m. We calculated the average peak frequency for the cluster channels, for each subject, and correlated those with the staircase and fitted SOA.

## Results

Participants performed a visual temporal discrimination task in which they were presented with two grids partially filled with annuli separated by SOAs of varying duration. If both grids were integrated, only one position was not filled with an annulus. Participants’ task was to report by button press whether they detected the empty position, i.e., whether they were able to temporally integrate both stimuli.

### Behavioural results

In a staircase procedure prior to the experiment, we determined the SOA, at which participants were able to detect the empty position in 50% of the trials (60.0 ± 7.7 ms; mean ± SEM). The average threshold SOA was 60 ms (± 38.45 ms, minimum 16 ms and maximum 144 ms). In the main experiment, we determined recognition rates as a function of different SOAs (short, threshold SOA, long, intermediate plus and minus SOAs, see Methods). For short SOA, recognition rates were at 0.80 ± 0.03 (mean ± SEM); for intermediate minus at 0.55 ± 0.04 (mean ± SEM), for threshold SOA at 0.45 ± 0.03 (mean ± SEM), intermediate plus at 0.42 ± 0.03 (mean ± SEM) and long SOA at 0.31± 0.04 (mean ± SEM) (Fig. 2). A one-way ANOVA revealed a significant main effect of SOAs (F = 20.9, p < 0.001; Fig. 2). Post hoc Tukey-Kramer tests showed that the short SOA differed significantly from all other SOAs (all pairwise comparisons p < 0.001). The intermediate minus SOA differed significantly from the intermediate plus SOA (p = 0.013) and the long SOA (p = 0.004). Post hoc tests showed no significant differences between intermediate minus SOA and threshold SOA (p = 0.212), threshold SOA and intermediate plus SOA (p = 0.769), threshold SOA and long SOA (p = 0.564), and intermediate plus SOA and long SOA (p = 0.999). Single subject behavioural data is shown in the supplementary material (Figure S1).

**Figure 2:**
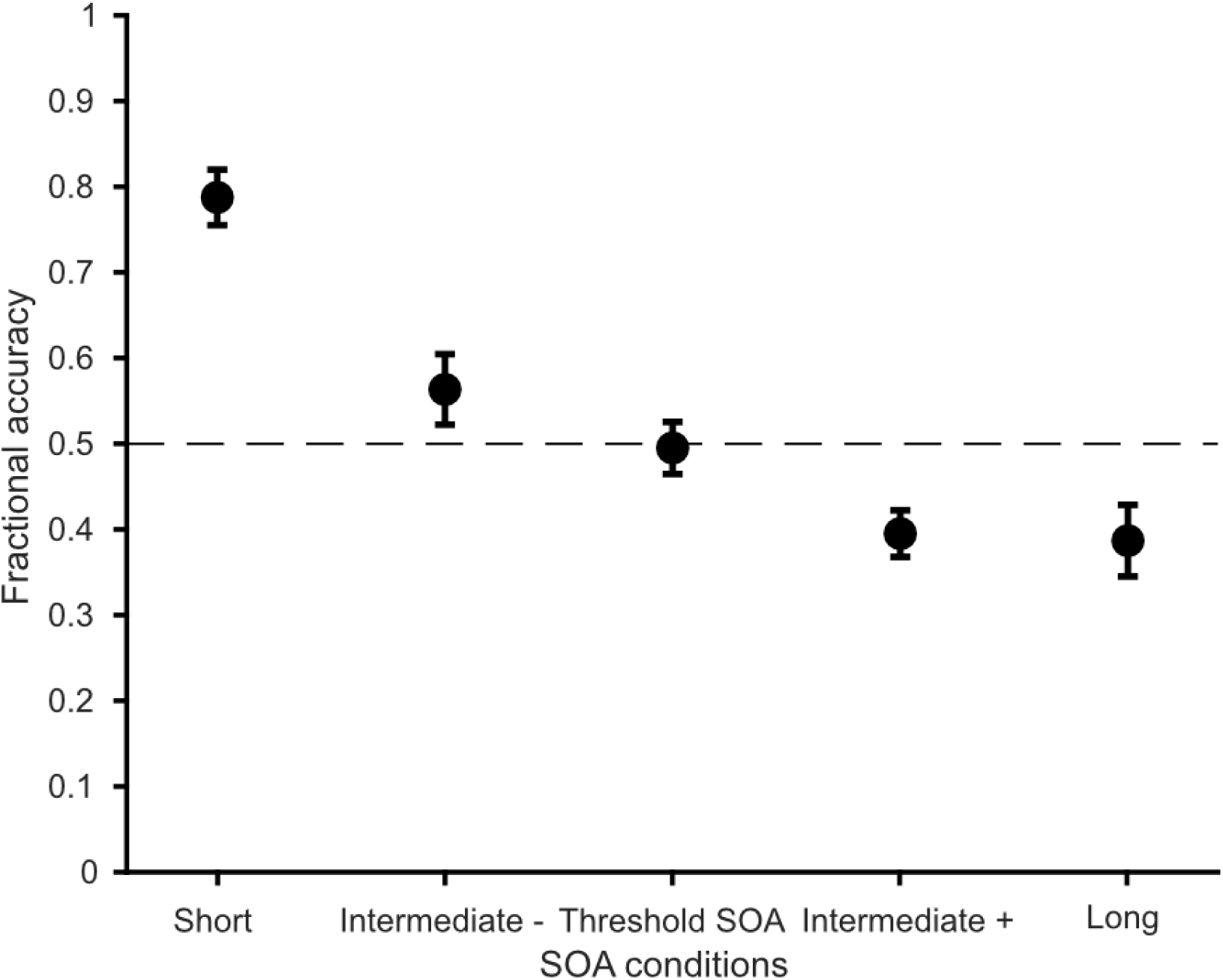
Behavioural performance. Fractional accuracy is plotted against the different SOA conditions. The dashed line marks the 50 % mark indicating the performance level we expected to see for the threshold SOA. Data are presented as mean ± SEM. A one-way ANOVA reveals a significant effect of SOA conditions (p < 0.001).

### Phase contrasts

To investigate whether perception is influenced by the phase of neuronal oscillations, we split all trials with a threshold SOA according to participants’ perception (i.e., correct, and incorrect trials). Next, we analysed putative phase differences between correct and incorrect trials by computing the phase opposition sum (POS, VanRullen, 2016) on source level for each time-frequency-grid sample and subsequently performing a cluster-based permutation test. The average trial count for correct trials was mean = 87.4 (± 34.343 STD) and for incorrect trials mean = 81.2 (± 25.146). The results revealed a significant cluster (p = 0.048, summed t-values: 2722.4, t-value range: [2.07 6.57]) between -0.8 s to - 0.5 s and between 6 Hz to 20 Hz, showing the most pronounced effect between 8 Hz and 16 Hz (Fig. 3B). The significant cluster was located at the *parietal lobe* (Fig. 3A left), and more prominent in the left hemisphere (Fig. 3A, right). We tested for systematic POS differences between high and low power trials, by recalculating the POS values for high and low power trials. We averaged the POS values per subject for the channel-time-frequency points we reported above. We found no power difference between POS values (t = -0.61, p = 0.577).

**Figure 3:**
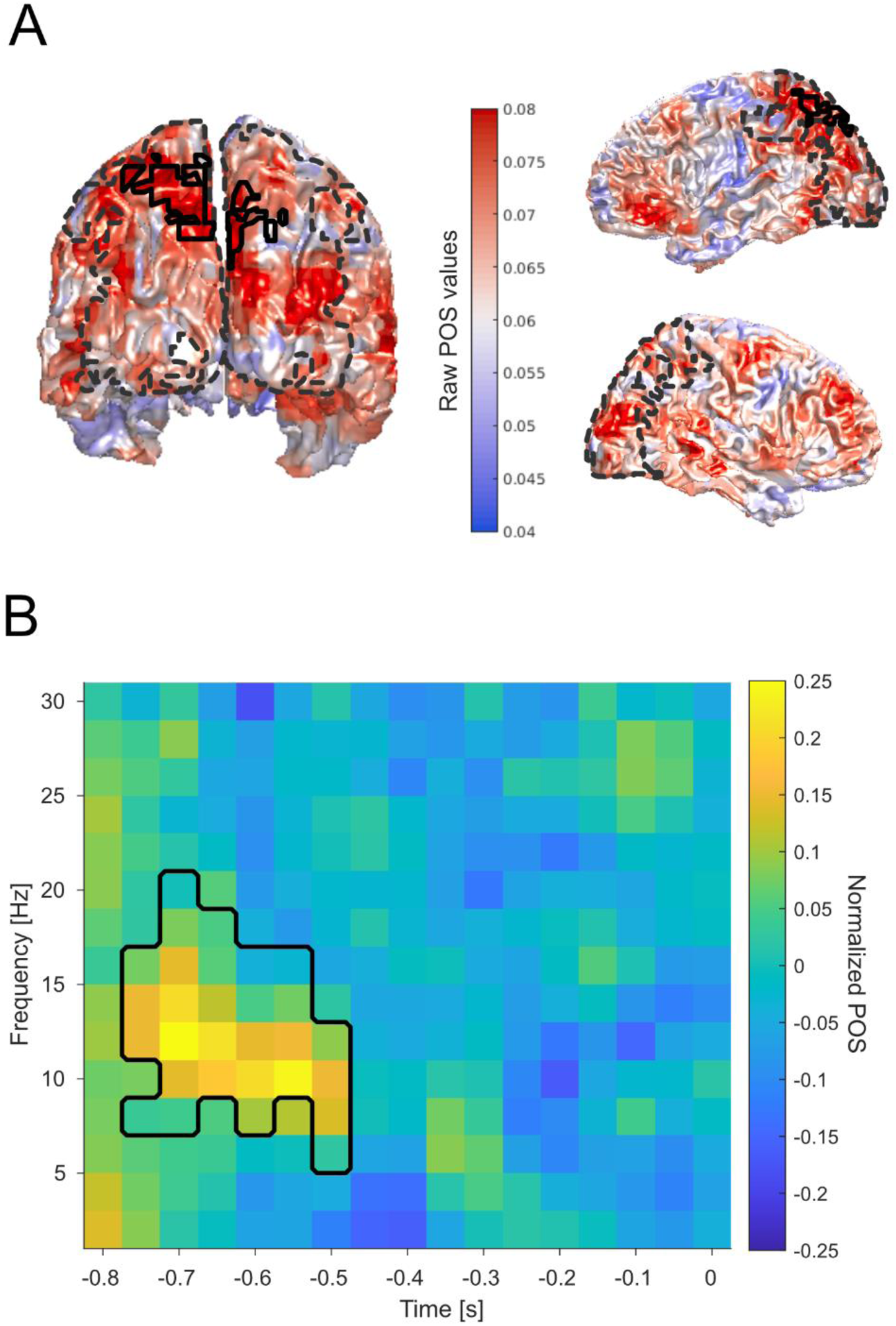
Phase opposition sum (POS) effects. A) POS values projected on a template MNI brain viewed from posterior (left) and right and left view (right). The black outline indicates the significant region/virtual channels (p = 0.048, summed t-values: 2722.4, t-value range: [2.07 6.57]. The dashed line indicates the region of interest for statistical comparison. B) Averaged normalized phase opposition values over all significant virtual channels of the cluster (see A). The black outline indicates the significant time-frequency range.

### Phase and behavioural performance

We investigated how behaviour was modulated by the momentary phase. To this end, we computed for each participant the phase for all trials at the time-frequency sample showing the highest POS value. Next, we binned single trials according to their momentary phase in six bins and averaged for each bin participants’ responses (Fig. 4). A one-way ANOVA excluding the phase bin containing the aligned highest responses (bin 0) revealed a significant effect of phase bin on accuracy (F = 3.9, p = 0.005). A post hoc Turkey-Kramer test showed a significant difference between bin π and bin 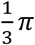 (p = 0.035) and between bin 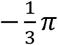 and bin 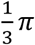 (p = 0.003), and a trend towards significance difference between bin 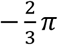 and 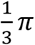 (p = 0.053). We additionally restricted the analysis to the alpha band (8-12 Hz), following the same pipeline but with a different frequency range of interest. The results showed no significant influence of phase on behaviour, when restricted to the alpha phase (F = 0.58, p = 0.681).

**Figure 4:**
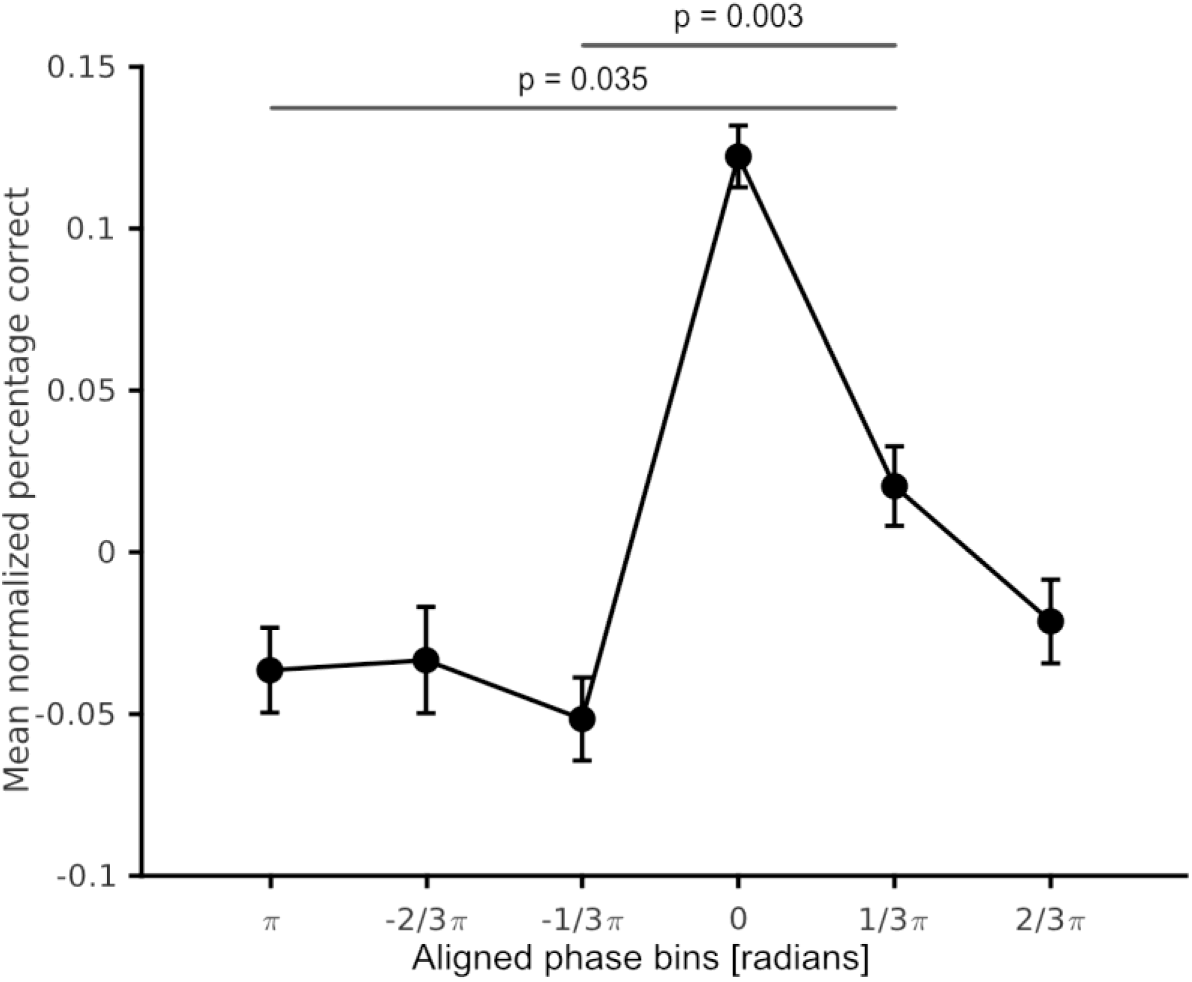
Phase dependence of behaviour. Normalized fractional accuracy plotted against the binned prestimulus phase. Data are shown as mean ± SEM. An ANOVA revealed a significant effect of phase bins (F = 3.87, p = 0.005). Black lines indicate significant differences between phase bins in post-hoc pairwise comparisons.

### Phase and ERF

We binned trials according to their momentary phase in equally spaced phase bins. Next, we computed per participant the ERF t values for all trials in each bin and averaged the ERF t values across participants. A one-way ANOVA showed a significant effect of phase bins (F = 3.4, p = 0.011; Fig. 5). A post hoc Turkey-Kramer test showed a significant difference between bin 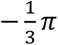 and bin 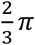 (p = 0.016) and a trend towards a significance between bin 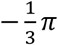 and bin 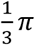 (p = 0.064). Additionally, we tested if these effects were also found, when restricted to the alpha band. The same analysis was performed but this time, choosing ERF values between 8 – 12 Hz and averaging over these frequency points. Our additional analysis showed that there was no difference between phase bins when restricted to the alpha band (F = 1.98, p = 0.102).

**Figure 5:**
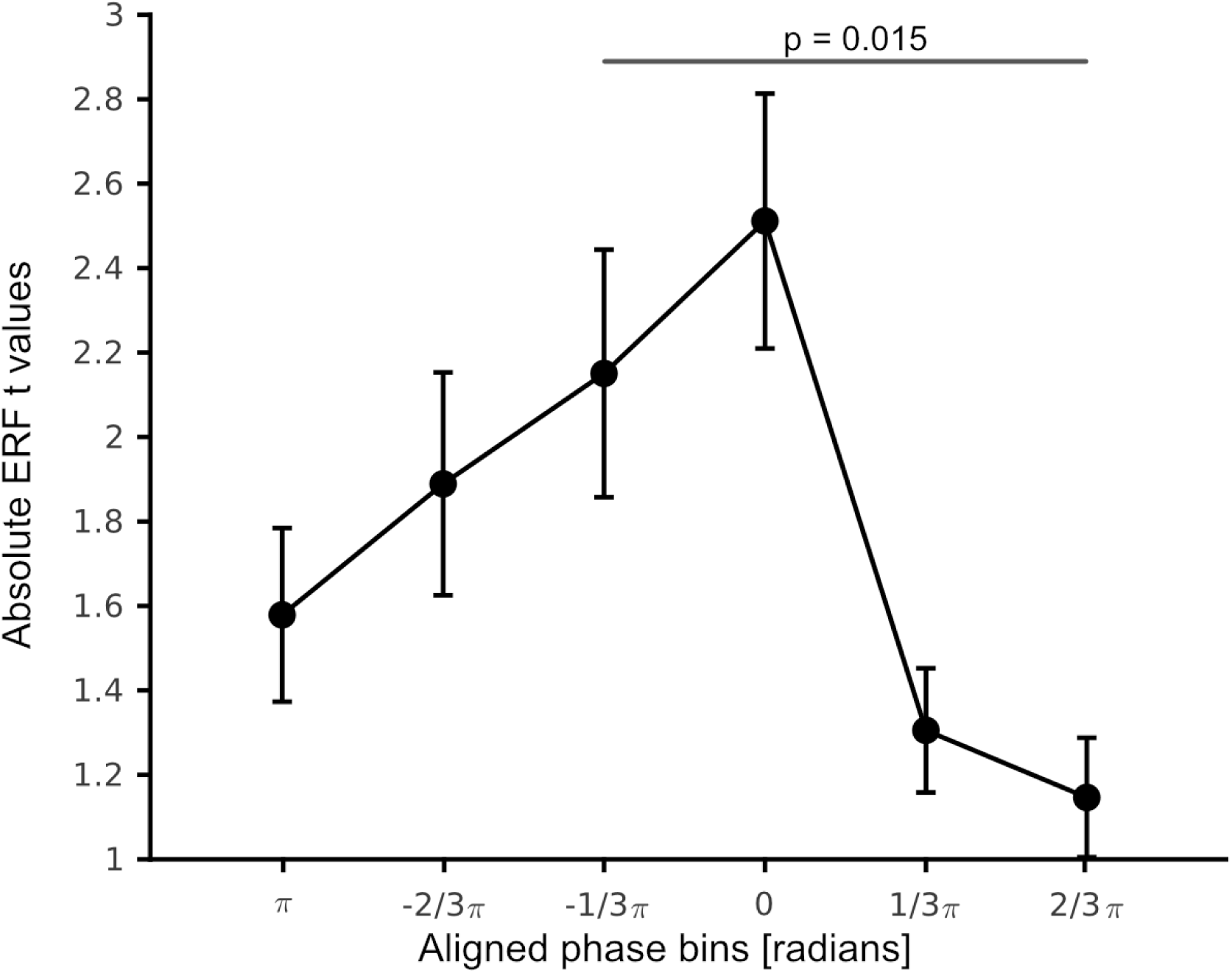
Phase dependence of ERF amplitude. ERF t values plotted against the binned pre stimulus phase. Data are shown as mean ± SEM. An ANOVA revealed a significant effect of phase bins (F = 3.4, p = .011). Black lines indicate significant differences between phase bins in post-hoc pairwise comparisons.

### Correlation of peak frequency and threshold SOA

We investigated whether the individual peak frequency correlates with individual threshold SOA. Therefore, we correlated peak frequency with the estimated SOA prior to the experiment (staircase) (Fig. 6A) and the fitted (Fig. 6B) threshold SOA (see Methods for details). For three participants the fitted threshold SOA could not be determined reliably (R^2^ < 0.33). These participants had to be excluded from the respective correlation analysis. We did not find any significant correlation between individual peak frequencies and staircase threshold SOAs (r = 0.25, p = 0.26) nor between peak frequencies and fitted threshold SOA (r = 0.18, p = 0.43). As for our supplementary analysis, where we correlated power peak frequency with the threshold and fitted threshold SOA, we found no significant correlation between either staircase (r = -0.1, p = 0.64) nor the fitted threshold SOA (r = -0.24, p = 0.24). Lastly, we restricted the frequency range of interest of our initial analysis to the alpha band 8-12 Hz, but because most peak were found initially int the alpha band, this restriction did not change the results. For the staircase threshold SOA (r = 0.25, p = 0.26) nor for the fitted threshold SOA (r = 0.18, p = 0.43) was a significant correlation found for the peak frequency.

**Figure 6:**
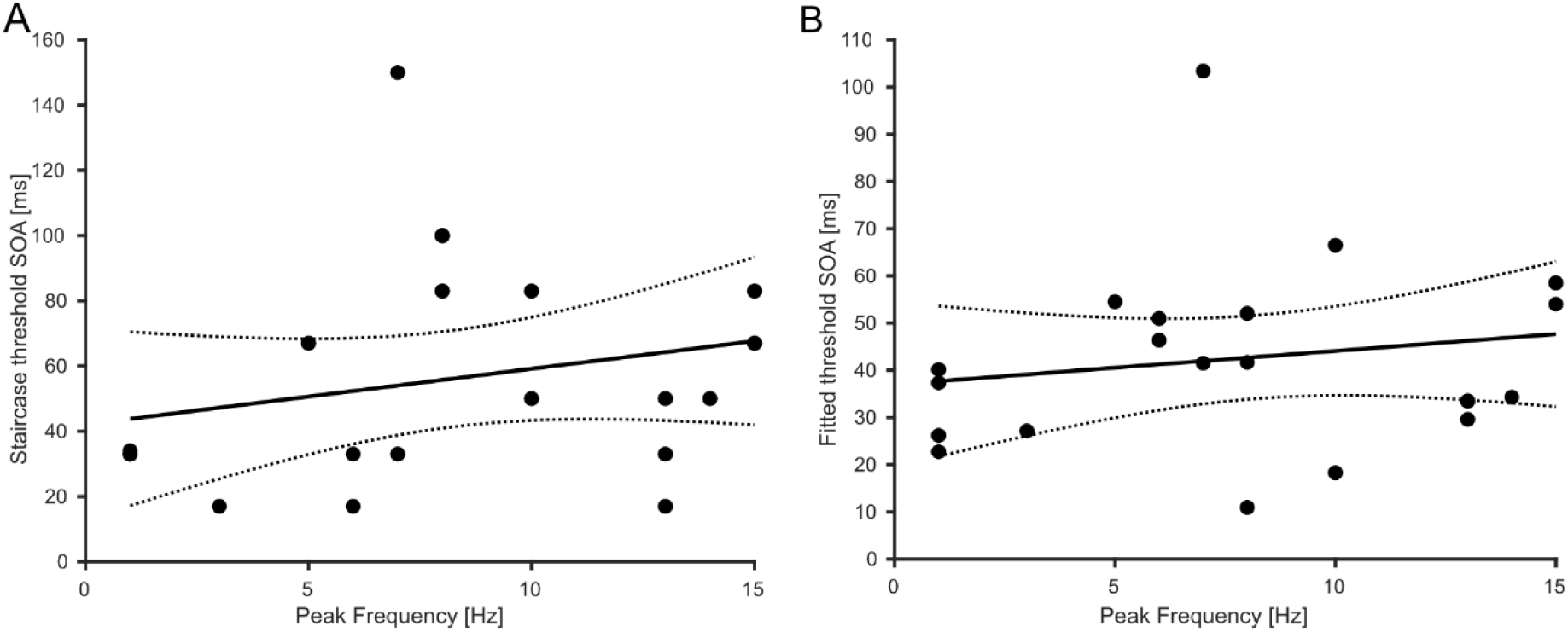
Correlation between peak frequency and threshold SOAs. A) Threshold SOA determined prior to the experiment plotted against the peak frequency (see Methods for details). Pearson correlation revealed no significant correlation (r = 0.25, p = 0.26). B) Same as A) but this time correlation between estimated fitted threshold SOA and peak frequency (see Methods for details). Pearson correlation revealed no significant correlation (r = 0.18, p = 0.423). The black line represents the linear regression, the dotted lines the 95% confidence intervals for the mean.

## Discussion

There is an ongoing debate whether prestimulus phase of neuronal oscillations influences perception (e.g., Keitel et al., 2022; Harris, 2023). Most evidence favouring a crucial role of phase stems from visual detection tasks, here, we investigated how prestimulus phase modulates visual temporal integration. We found phase differences when comparing correct with incorrect trials. These phase effects were in the frequency range between 6-20 Hz, and - 0.8 to -0.5 s prior to stimulus presentation, mainly in parieto-occipital areas. Additionally, accuracy decreased when deviating from the preferred prestimulus phase. Finally, prestimulus phase modulates poststimulus visual processing as the N170 amplitude attenuated when deviating from the individually preferred phase. However, we found no evidence that the individual frequency of the prestimulus phase correlates with the temporal integration threshold.

Previous studies investigating visual detection tasks reported phase effects in the alpha-band (Busch et al., 2009; Mathewson et al., 2009; Landau and Fries, 2012; Kraut and Albrecht, 2022; Zazio et al., 2022), in our study the effects are largest in the range 8-16 Hz, but also exceeding to the low beta-band (20 Hz). These effects could be due to inter-individual differences or is a true broadband effect. Our analysis is not suited to investigate this effect further. However, the higher frequencies in our study might depend on task demands. It has been reported that alpha peak frequency shifts to higher frequency in cognitively more demanding tasks (Haegens et al., 2014). Similarly, studies have shown that occipital alpha peak frequencies differ between visual integration or segregation task (Wutz et al., 2018; Sharp et al., 2022; Han et al., 2023). Relevant frequency shifted with task difficulty also in a visual search task (Merholz et al., 2022). In line with these findings, the higher frequencies here reported might be explained by higher task difficulty of a temporal discrimination task compared to detection tasks. On the other hand, Wutz et al. found the opposite effect that alpha frequency decreased in their integration task compared to their segregation task (Wutz et al., 2018). A crucial difference in study design is, that participant had to change strategies during the experiment. It could be that due to the switch the difference in alpha peak frequency occurred. While most study expected effects or tested effect over the occipital lobe (Wutz et al., 2018; Alexander et al., 2020; Fakche et al., 2022; Zazio et al., 2022). We found also effects over the parietal lobe. One possible explanation could be that our source reconstruction is imprecise. Another could be that higher order visual areas are involved in solving this task. When considering the two-pathway hypothesis, which states that the “where” pathway end in the parietal lobe, it would be possible that these areas are also activated in a task where a target must be spatial located (Mishkin et al., 1983).

The reported performance effects (decreasing when deviating from preferred phase) are smaller compared to other studies (∼7% difference between highest and lowest bin compared to ∼13%, Busch et al., 2009; Dugue et al., 2011; Baumgarten et al., 2015; Fakche et al., 2022). Task difficulty could be a leading cause for the difference, while the effect pattern is comparable. In addition, we found that prestimulus phase modulates the amplitude of the poststimulus N170 component. In line with our finding, prestimulus phase has been reported to modulate global field potential (Busch and VanRullen, 2010; Dou et al., 2022; note that Dou et al. found these effects only for high power trials), TMS-evoked ERPs (Romei et al., 2012; Fakche et al., 2022), and visual awareness negativity (Krasich et al., 2022). In sum, these studies suggest across a variety of different tasks that prestimulus phase can modulate postimulus components.

While our results are comparable to detection task results, the underlying mechanism is not well understood. Earlier studies proposed the idea that stimulus presentation triggers a phase shifted towards a preferred phase, inducing a phase reset (Makeig et al., 2002). When prestimulus phase is at the preferred phase, stimulus processing is optimal, leading to increased evoked responses and improved performance. Whereas, when phase has to shift toward the preferred phase, processing is less optimal, leading to decreased evoked responses and behaviour. This could be investigated by comparing phase angle values before and after phase reset. Such analyses, however, might be practically less straightforward due to potential temporal smearing effects by the methods for phase analysis (Brüers and VanRullen, 2017).

Neuronal phase might also indicate the current excitability of the underlying neuronal population (Bishop, 1932; Lakatos et al., 2007; Fakche et al., 2022). In theory, cortex excitability changes rhythmically with phase, while excitability is highest at a preferred phase (Mazaheri and Jensen, 2006; Jensen and Mazaheri, 2010; Schaworonkow et al., 2019). Neuronal processing would be facilitated at such a preferred phase and putatively lead to higher ERPs. Stimulus processing would be more effective and putatively lead to better behavioural performance. Cortical excitability has also been repeatedly linked to the power of alpha oscillations (e.g. Romei et al., 2008; Lange et al., 2013; Dou et al., 2022). Here, low alpha power is beneficial for perception, while high alpha power suppresses excitability. Notably, phase effects on behaviour and neuronal processing have often been reported for high but not for low alpha power trials (Mathewson et al., 2009; Alexander et al., 2020; Dou et al., 2022; Fakche et al., 2022; Ozdemir et al., 2022). The reason could be, when power is low, cortical excitability is high and phase effects are less notably or irrelevant. In contrast, when power is high, excitability is low and phase matters more strongly as excitability differs more strongly between different phases.

Between studies, there is a high variability regarding the latency of the reported phase effects. Some report effects close to stimulus onset (∼ 100-200 ms before stimulus onset; e.g., Alexander et al., 2020; Fakche et al., 2022; Zazio et al., 2022) other studies – including the present study - report phase effects several 100 ms before stimulus onset (Hanslmayr et al., 2013; Baumgarten et al., 2015; Kraut and Albrecht, 2022). We can only speculate about the causes of these discrepancies. One reason might be that the timing of effects depends on the task (see above discussion on the frequency band). Temporal smearing effects of peri- or poststimulus effects into the prestimulus period (Brüers and VanRullen, 2017) might also overshadow putative phase effects close to stimulus presentation. Another explanation could be that slow frequencies orchestrate faster frequency. Evidence stems from reported power-phase coupling between higher and lower frequencies (Palva, 2005; Canolty et al., 2006; Axmacher et al., 2010; Voytek, 2010). Similarly, studies have reported phase-phase-coupling of neuronal oscillations in different frequency bands (Belluscio et al., 2012; Scheffer-Teixeira and Tort, 2016). Underlying (ultra-)slow oscillation orchestrating alpha activity on a network level would lead to a rhythmic occurrence of phase effects, with a visible component several ms before stimulus presentation. Respiration could be such a slow oscillation, by increasing the overall oxygen level, which would lead to higher excitability. Respiration has an effect on behavioural measures (Johannknecht and Kayser, 2022) and modulates alpha power at rest and during detection tasks (Kluger et al., 2021a; Kluger et al., 2021b). In addition, the slow cardiac cycle influences neuronal activity and behavioural performance rhythmically (Kim et al., 2019; Al et al., 2020). Such slow fluctuations have also been reported for perception (e.g., Monto et al., 2008; Palva and Palva, 2012). By investigating longer prestimulus periods of several seconds, future studies might shed more light on such rhythmic patterns and how they affect phase and other components of neuronal oscillations.

Finally, we investigated if a specific frequency band is relevant for unimodal visual temporal integration. Recent studies found evidence for a correlation between individual alpha peak frequencies and behavioural performance in visual or multimodal studies (Samaha and Postle, 2015; Keil and Senkowski, 2017; Minami and Amano, 2017; Bastiaansen et al., 2020; Migliorati et al., 2020; Venskus and Hughes, 2021; Venskus et al., 2021). A common interpretation of such correlations is that the cycle of a neuronal oscillation might reflect temporal integration windows, while the frequency shapes the window size. Following a slightly different approach, by taking the frequency with the highest individual phase effect (Baumgarten et al., 2015), we found no evidence for an influence of frequencies on perception or temporal integration.

In light with ongoing debate if our perception is a rhythmic or discrete process, as described by van Rullen (VanRullen and Koch, 2003), we found support for the rhythmic hypothesis. In short, the idea of rhythmic perception is that the intensity or probability of perception changes with phase. While the idea of discrete perception is that, the timing of perception is modulated by the underling oscillation creating a window of integration. Our results showed that phase correlates with behaviour and individual participants seem to have a preferred phase where perception is high, while we found no support for an integration window. There we suggest that unimodal visual temporal integration is a rhythmic process, in a wider frequency range as expected, and potentially even modulated by even slower underlying oscillations.

A limitation of this study is that the estimation of individual phase effects is typically less stable as determining peak frequencies. Such noisy estimation might influence the results. In addition, we observed high subject variability in the behavioural data and phase opposition effects (absolute value, timing, frequency). Our analysis captures the overlapping effects on group level. Other study designs would be more suited to capture individual differences.

In addition, it should be noted that several studies report nullfindings with respect to phase effects on perception or neuronal activity (Ruzzoli et al., 2019; Michail et al., 2022; Zazio et al., 2022; Melcón et al., 2023). Here, we can only speculate about the discrepancies. In our study, phase effects were very local in space (see also Palva, 2005; Baumgarten et al., 2015). In addition, typically phase effects are smaller than power effects. Therefore, phase effects might be too small to be detected or overshadowed by other effects, especially when analysed on sensor level, as spatial leakage of other effects might be stronger than the actual phase effect. It also seems that phase effects are only visible in tasks near the perceptual threshold, but not for easier tasks. It remains to be studied whether phase plays a continuous role in our perception or is only relevant near the perceptual threshold. Finally, potential phase effects might be overlooked due to methodological problems in the analyses (Harris, 2023).

In conclusion, we found an effect of phase of alpha oscillations in a visual temporal discrimination task and on neuronal processing. These results suggest that perception is a rhythmic process and/or that the phase modulates stimulus processing. While these results are of correlative nature, future studies might investigate causal relationships between phase and perception/neuronal processing. To test a putative causal relationship, phase resets could be triggered externally and the consequences on behaviour can be tested.

## Author Contributions

JL designed research; MJ performed research; MJ and JL analysed data; MJ, JL and AS wrote the paper.

## Acknowledgment

We thank the German Research Foundation for funding this project (Grand LA 2400/8-1). Computational infrastructure and support were provided by the Centre for Information and Media Technology at Heinrich Heine University Düsseldorf. We thank the MRI Core Facility of the Medical Faculty of Heinrich Heine University for the support with MRI measurements.

## Conflict of interest

The authors declare not conflict of interest.

## Founding source

German Research Foundation (Grand LA 2400/8-1).

## Appendix

## Supplementary Material

**Figure S1:**
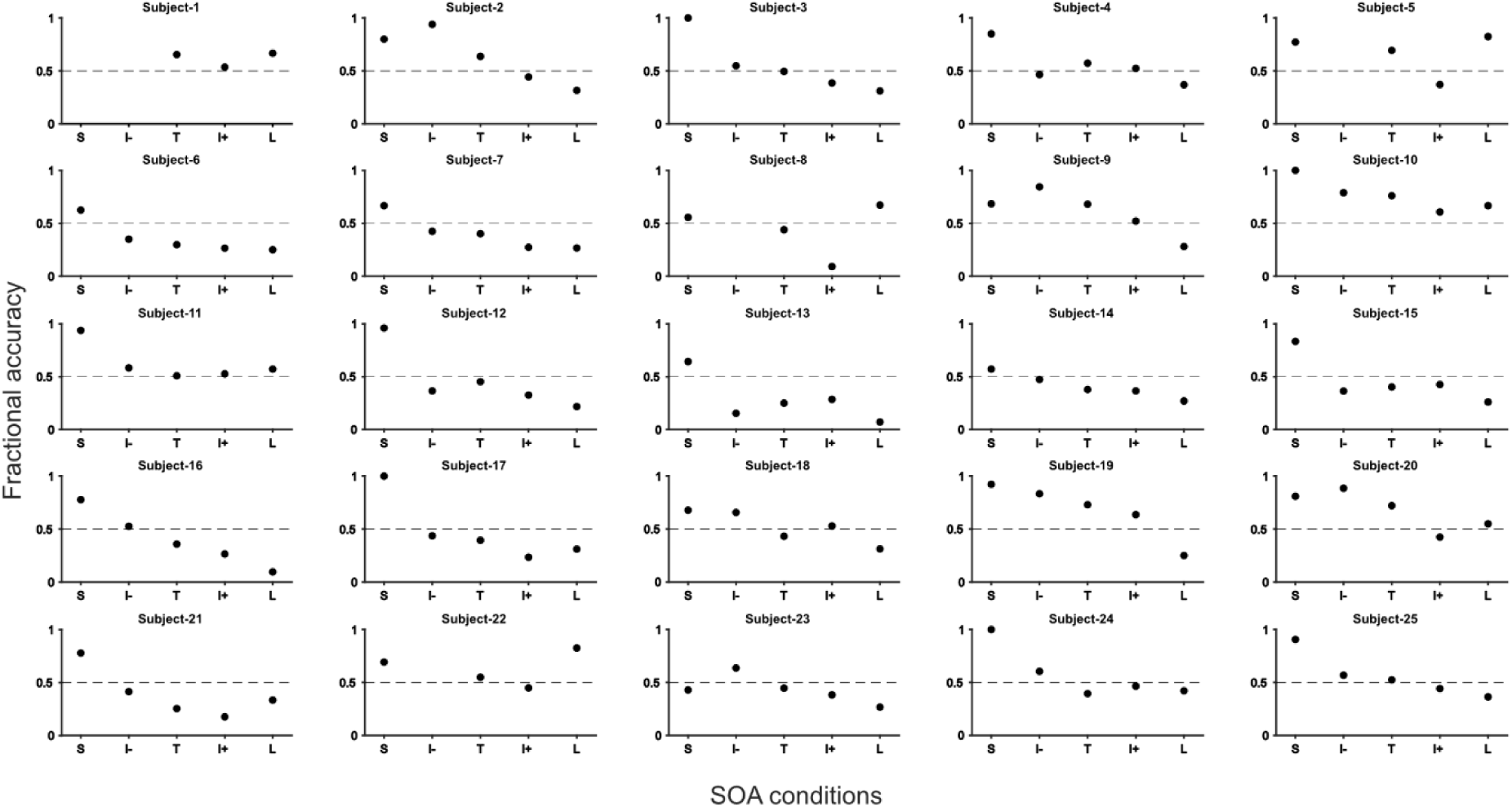
Behavioural performance on single subject level. Fractional accuracy is plotted against the different SOA conditions. The dashed line marks the 50 % mark indicating the performance level we expected to see for the threshold SOA. Conditions are ladled as S for Short, I- for Intermediate-, T for Threshold, I+ for Intermediate+ and L for long. Some conditions are missing for some participant either because the Threshold SOA was the shortest SOA, or because the intermediate- and short SOA condition were the same. If the later was the cast, both conditions were combined to the short condition.

**Table S1:** Statistical Test Overview. The table shows for which data structure, we preformed which test and the confidence intervals.

| Data Structure | Type of test | Power |
| --- | --- | --- |
| Behavioural Data (non-normal distributed) | One-way ANOVA | CI 95 % |
| Phase opposition sum values, normalised (normal distribution) | Cluster based permutation testing | CI 95 % |
| Phase binned behavioural data (non-normal distributed) | One-way repeated ANOVA | CL 95% |
| Phase binned ERP data (non-normal distributed) | One-way repeated ANOVA | CL 95% |
| Peak frequency values (non-normal distributed) | Pearson Correlation | CL 95% |

## Notes

### Competing Interest Statement

The authors have declared no competing interest.

